# Reduced protein expression in a virus attenuated by codon deoptimization

**DOI:** 10.1101/106799

**Authors:** Benjamin R. Jack, Daniel R. Boutz, Matthew L. Paff, Bartram L. Smith, James J. Bull, Claus O. Wilke

## Abstract

The engineering of hundreds of synonymous codon changes into a viral genome appears to provide a general means of achieving attenuation. The mechanistic underpinnings of this approach remain enignmatic, however. Using quantitative proteomics and RNA sequencing, we explore the molecular basis of attenuation in a strain of bacteriophage T7 whose major capsid gene was engineered to carry 182 suboptimal codons. As expected, there was no evident effect of the recoding on transcription. Proteomic observations revealed that translation is halved for the recoded major capsid gene, and a smaller reduction applies to a few genes downstream, potentially caused by translational coupling. Viral burst size is also approximately halved, and the fitness drop accompanying attenuation is compatible with the reduced burst size. Overall, the fitness effect and molecular basis of attenuation by codon deoptimization are compatible with a relatively simple model of reduced translation of a few genes and a consequent diminished virion assembly. This mechanism is simpler than that operating in eukaryotic viruses.

## Introduction

A recent methodological advance in the development of attenuated viral vaccines has been the design of genomes with hundreds of synonymous codon changes that are collectively suboptimal (Burns et al, 2006; Mueller et al, 2006; Coleman et al, 2008; Burns et al, 2009; Wimmer et al, 2009; Mueller et al, 2010; Bull et al, 2012; Nougairede et al, 2013; Shen et al, 2015). The nature of the design varies from the replacement of common codons with rare codons to merely shuffling existing codons to create uncommon codon pairs. By maintaining wild-type protein sequences, the recoding method retains the antigenic profile of the wild type while reducing viral growth rate and virulence. Based on the premise that silent codon changes have individually small effects, the method should not only allow the tunable crippling of viral growth to arbitrary degree but also profoundly retard the re-evolution of high viral fitness. Both predictions have been supported empirically. Thus, poliovirus, influenza, several arboviruses, and a bacteriophage all exhibit quantitative fitness declines after synonymous codon changes (Burns et al, 2006; Mueller et al, 2006; Coleman et al, 2008; Mueller et al, 2010; Nougairede et al, 2013; Shen et al, 2015; Bull et al, 2012), and fitness recovery during viral growth over hundreds of generations is at best slow (Burns et al, 2006; Coleman et al, 2008; Bull et al, 2012; Nougairede et al, 2013).

The broad success of attenuation from synonymous codon changes in different viruses and with different designs could arise from a common underlying mechanism. Yet, there is ongoing debate about how synonymous codon changes affect fitness and thus what that mechanism could be. One popular hypothesis is that codon usage controls translational efficiency, in turn affecting the rate of protein synthesis (Fredrick and Ibba, 2010; Plotkin and Kudla, 2010; Tuller et al, 2010; Shah and Gilchrist, 2011; Zur and Tuller, 2016). Under this hypothesis, highly expressed and functionally important genes are encoded by optimal codons to increase translational efficiency. However, a simple codon ‘optimality’ model cannot explain the attenuation attained by merely shuffling codons—since the abundance of the different codons is not being changed by shuffling, the suboptimality must be due to something besides codon abundance. Likewise, some highly-expressed genes in cyanobacteria and Neurospora have non-optimal encodings (Xu et al, 2013; Zhou et al, 2013). In other cases, codon usage determines expression through transcription, not translation (Zhou et al, 2016). In the face of so many seemingly contradictory observations, further advances in this research program would benefit from the identification of a common mechanism for viral attenuation, or at least benefit from the demonstration that different mechanisms are involved.

Resolving the basis of attenuation in viral systems is partly hampered by the sequence of life-history steps between the initial effects of codon changes and the final emergence of assembled virions. The initial impact may lie in transcription or translation of one or more genes, but the effect on the number of virions will depend on which proteins are limiting during assembly. The same modification of an early life-history stage may have different fitness effects in different viruses. Ultimately, it may be necessary to interpret the genome engineering in the context of a system-wide, comprehensive model of the viral life cycle. Such is our motivation.

In the bacterial virus T7, recoding the major capsid protein (gene *10A*) with synonymous codons reduced the fitness of the phage (Bull et al, 2012). The engineering of T7 substituted codons that were underutilized in *E. coli* (the T7 host) in place of codons that were highly utilized in the host. The major capsid protein, which forms the head of the T7 phage particle, is the most abundant and highly-expressed phage protein (Dunn and Studier, 1983). In the phage genome with the most extensive set of gene 10A synonymous codon replacements, the fitness was 35.7 doublings per hour compared to 43.2 doublings per hour in the wild type (Bull et al, 2012). This difference translates to a 180-fold decline in descendants produced per hour. Fully 182 codons were changed, just over half the number of codons in the major capsid gene; in addition, the first 20 codons were not altered to avoid disrupting translation initiation processes.

After evolving the recoded phage for 1000 generations, fitness recovered to about half the difference from the wild-type phage (on a log scale). Thus, recoding gene *10A* induced a moderately stable fitness reduction. A mere 9 nucleotide changes were responsible for the fitness recovery, and 7 fell outside the recoded region, shedding little light on the underlying mechanism of fitness reduction.

Here, we apply new methods to continue exploration of the T7 attenuation. Our purpose is to develop a comprehensive model of the way silent codon changes cause reduced fitness. As part of this effort, we propose and test three mechanistic models that could explain the fitness reduction in recoded T7. In testing those models, we apply proteomic methods, RNA sequencing, and various phenotypic measures in a systems approach to understanding the basis of attenuation. We find that recoding gene *10A* reduces protein abundances of gene *10* and also of several downstream genes. From there, we address the impact of protein abundances on viral fitness components (burst size and lysis time), ultimately connecting these measurements to a model that describes actual fitness.

## Results

### Codon deoptimization reduces capsid protein abundances

As codon deoptimization is thought to affect translational efficiency, we propose three models in which recoding gene *10* (capsid protein) affects protein abundances in T7. In model 1, codon deoptimization slows translation of gene *10* and reduces the abundance of the capsid protein only. In model 2, deoptimization depletes the ribosome pool by creating high ribosomal densities on capsid protein transcripts, thus reducing translation of all viral proteins late in the infection cycle (Vind et al, 1993; Birch et al, 2012; Raveh et al, 2016). In model 3, codon deoptimization has intermediate effects between models 1 and 2: translation is impaired for gene *10* and downstream genes but not for upstream genes. The expectation in model 3 arises because T7 produces many polycistronic transcripts, leading to translational coupling of gene *10* and genes immediately downstream. Translational coupling has been observed in bacterial operons but has not been considered in the context of codon deoptimization (Oppenheim and Yanofsky, 1980; Schümperli et al, 1982; Aksoy et al, 1984; Tian and Salis, 2015). All three models assume that deoptimizing gene *10* will reduce the abundance of at least the capsid protein.

To differentiate between our three proposed models, we compared the T7 proteome during infection among wild-type, attenuated, and evolved phages using mass spectrometry-based protein quantitation. T7 is thought to encode 58–60 proteins, but only 19 are essential, and many have no known function (Dunn and Studier, 1983; Molineux, 2005) (e.g., some are thought to be homing endonucleases, selfish elements). All genes are encoded on the same strand, and expression order is linear; genes are numbered in order with essential genes having integral numbers (*1 -19*) and non-essential genes having fractional numbers. The genome is divided into 3 expression groups. Class I genes are the first to enter and are expressed from promoters at the entering end of the genome, transcribed by the host RNA polymerase (RNAP). The phage RNAP gene (numbered gene *1*) is the last of the class I genes and the first essential gene. All other genes are expressed from phage promoters, but nearly all transcripts are polycistronic, as there is only one terminator for phage RNAP (immediately after gene *10*), and there are only 17 phage promoters (Fig 8). Gene *1.1* is the first class II gene (Dunn and Studier, 1983). [Note that genes *1.1 –1.3* are sometimes also considered to be part of class I, because they are transcribed by both *E. coli* and T7 RNAP (Molineux, 2005).] Gene 6.5 is the first class III gene.

Because the wild-type phage lyses the cell at approximately 11 minutes after infection (Heineman and Bull, 2007; Bull et al, 2011) the proteome of infected hosts was sampled at 1, 5, and 9 minutes after phage addition to the culture; infection of cells is neither immediate nor synchronous upon phage addition to the culture, so these times are approximate post-infection values. By 9 minutes after infection, roughly 50 of the known or predicted T7 proteins were detected. All samples recovered approximately 4000 *E. coli* proteins. Since proteins have a much longer half-life than transcripts, and T7 has no known mechanism of degrading *E. coli* proteins (Molineux, 2005), we assumed that *E. coli* protein abundances remained constant over the 9 minute infection, and normalized the phage protein abundances to that of *E. coli* (Houser et al, 2015). Thus, we report all phage protein abundances as a proportion of *E. coli* protein content.

Abundances of the major capsid protein (a product of gene *10*) were of primary interest, as gene *10* is the most highly-expressed phage gene and is also the one deoptimized. Gene *10* comprises two protein products: the major capsid protein (*10A*) and the much less abundant minor capsid protein (*10B*). The minor capsid protein is produced after a frameshift and stop codon read-through of *10A* and, except for the C-terminal ~53 amino acids, is identical in sequence to the major capsid protein. Thus, our proteomic methods have limited ability to distinguish between the two protein products, so we combine abundance estimates into a single capsid protein measurement (see Materials and Methods). By 9 minutes after infection, capsid protein abundances in the attenuated strain were about half of those in the wild type (*p* < 0.05, paired *t* test, Fig 1). The capsid protein abundance for the evolved strain was intermediate.

**Figure 1:**
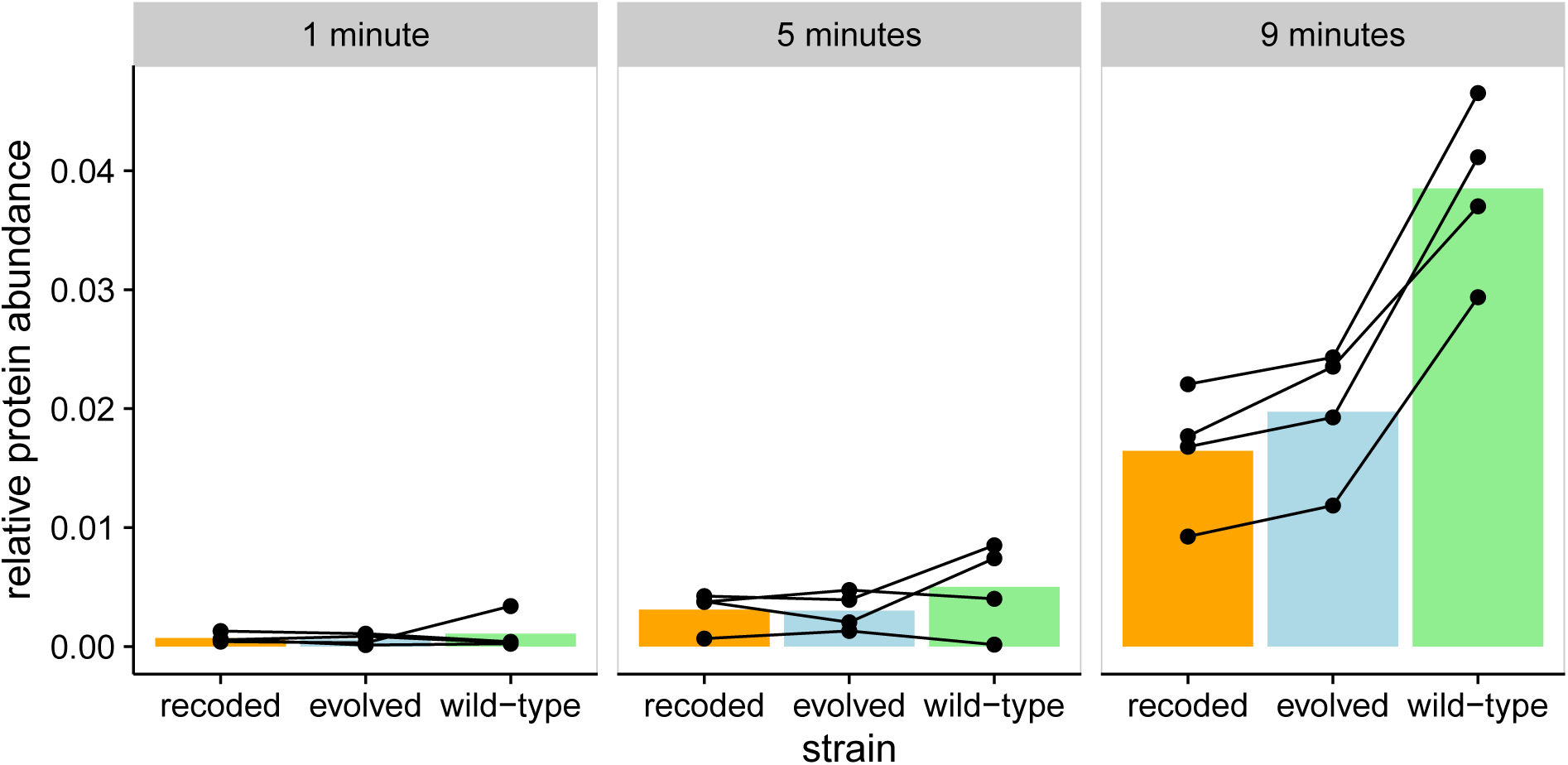
Recoding gene *10A* reduces capsid protein abundances. We measured protein abundance at 1, 5, and 9 minutes after infection. In the recoded (attenuated) strain, protein abundance for capsid protein after 9 minutes of infection is half of that of the wild type (*p* < 0.05, paired *t* test). The evolved strain also has significantly lower levels of capsid protein after 9 minutes. Each point represents a single measurement, and lines connect biological replicates.

Recoding of gene *10* could have reduced capsid protein abundance by reducing rates of translation elongation, thereby increasing the likelihood of ribosome stalling and fall-off. If ribosome fall-off were the dominant mechanism by which protein abundance was reduced, attenuation should have been accompanied by an excess of short peptides from the N-terminal end. Alternatively, if translation was slowed down without ribosome fall-off, a uniform distribution of peptides should be observed across the capsid protein. When mapping individual peptides recovered from the mass spectrometry proteomics, no systematic change was observed in the distribution of peptides across the protein (Fig 2). Thus, the recoded phage strains produced complete capsid protein, but in smaller quantities than that of the wild type.

**Figure 2:**
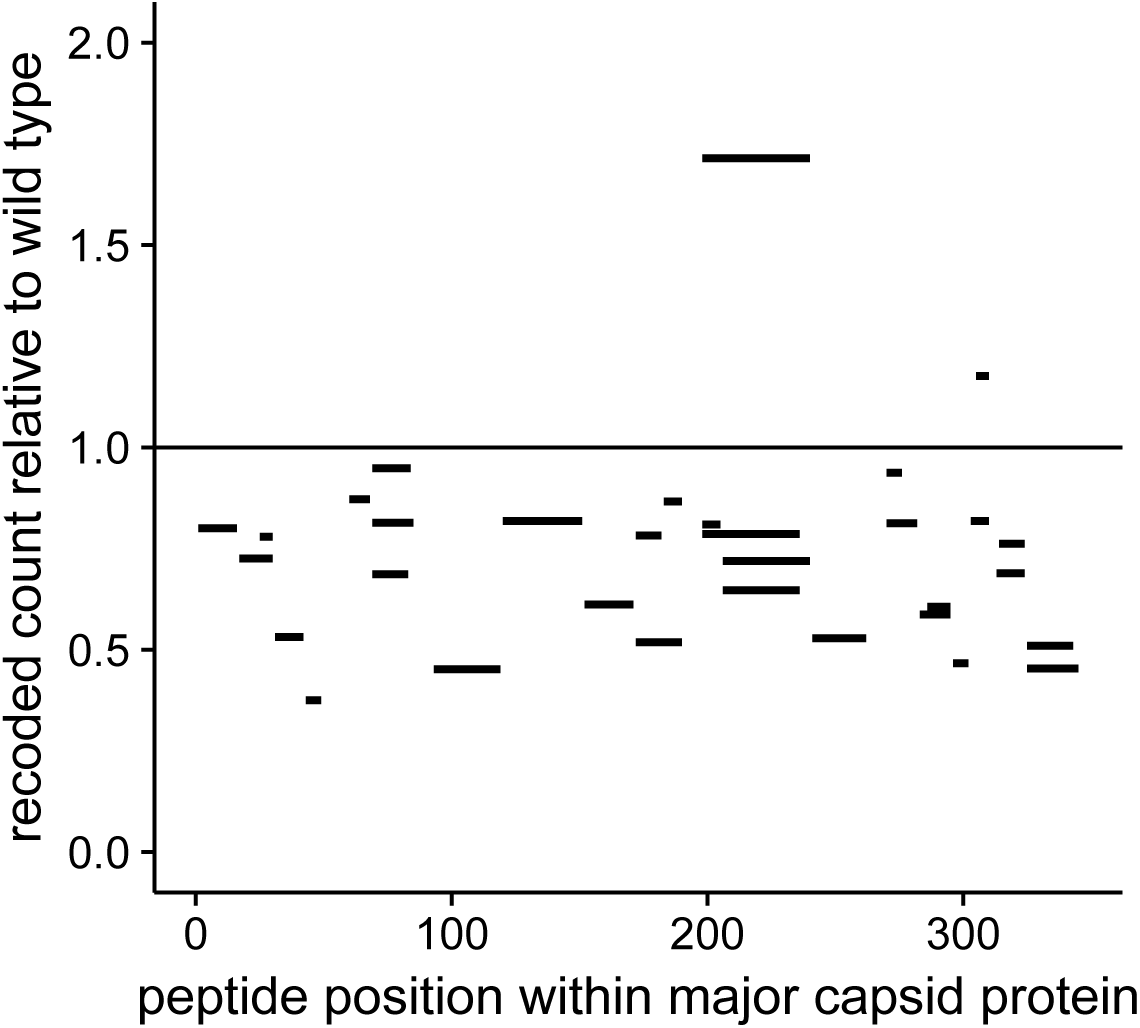
Peptides abundances are uniformly distributed across the capsid protein in the recoded T7 strain. Here we show the abundance of individual peptides within the capsid protein of the recoded strain, relative to the wild type. Data are from 9 minutes after infection. If translation consistently terminated before the stop codon in gene *10* due to recoding, we would expect the relative peptide abundance to systematically vary with the location of the peptide. Therefore, we tested a model that includes an interaction between strain and peptide location, and a model that includes no interaction (see Materials and Methods). We found that a model in which an interaction is included between the strain and peptide location fits the peptide count data no better than a model without interaction (log-likelihood = −1682.2, log-likelihood = −1682.4, respectively). Thus, in the recoded strain, there is no evidence of early translation termination.

### Codon deoptimization reduced some other class III protein abundances

If recoding a highly expressed gene saturates the pool of ribosomes by slowing translation of the recoded gene, the rate of translation of all T7 genes could ultimately decline (model 2, above). Models of T7 replication in *E. coli*, and limited experimental data, are consistent with protein synthesis being the rate-limiting step of T7 replication (although the evidence is at best weak and indirect) (Endy et al, 1997; You et al, 2002). Moreover, depletion of free ribosomes is common in *E. coli* transformed with highly expressed heterologous genes (Vind et al, 1993; Scott et al, 2010; Reuveni et al, 2011; Raveh et al, 2016).

The data allow us to measure other T7 protein abundances over time. Nine minutes post-infection, abundances of class III proteins p11, p12, p13, p14, and p15 in the recoded strain were lower than that of the wild type (false discovery rate FDR < 0.1, FDR-corrected *t* test) (Fig 3, Table 1). These proteins are mostly or all structural: tail tubular proteins (p11 and p12), and two internal virion proteins (p14 and p15), and a protein of unknown function but required for incorporation of other essential proteins in the virion (p13). The T7 endonuclease (p3), encoded by a class II gene, also showed reduced abundance in the recoded strain. Abundances of these class III proteins in the evolved strain again fall somewhere between that of the wild-type and recoded strains. Together, these results demonstrate that recoding gene *10* reduced the protein abundances of the capsid protein and several proteins encoded downstream of the capsid protein. The false discovery threshold we employed allows for one or two of those downstream proteins identified to be false positives, but the majority are likely real.

**Table 1:** Proteins in which abundance differs significantly (FDR < 0.1, FDR-corrected paired *t* test) between the wild-type and recoded strains of T7. Most differentially expressed genes are class III genes, with the exception of p3, a class I gene. The mean differences in abundance, unadjusted p-values, and false discovery rates are shown. Since eight genes fall below a false discovery rate of 0.1, we expect approximately one of these genes to be a false positive.

| protein | mean difference | $p$-value | false discovery rate |
| --- | --- | --- | --- |
| p3 | 2.7e-04 | 0.00014 | 0.0061 |
| p10 | 2.2e-02 | 0.00051 | 0.0110 |
| p13 | 8.4e-05 | 0.00180 | 0.0260 |
| p12 | 3.2e-04 | 0.00590 | 0.0510 |
| p14 | 2.3e-04 | 0.00560 | 0.0510 |
| p11 | 3.3e-04 | 0.00740 | 0.0530 |
| p15 | 2.9e-04 | 0.01400 | 0.0880 |
| p18 | 3.2e-04 | 0.01600 | 0.0880 |

**Figure 3:**
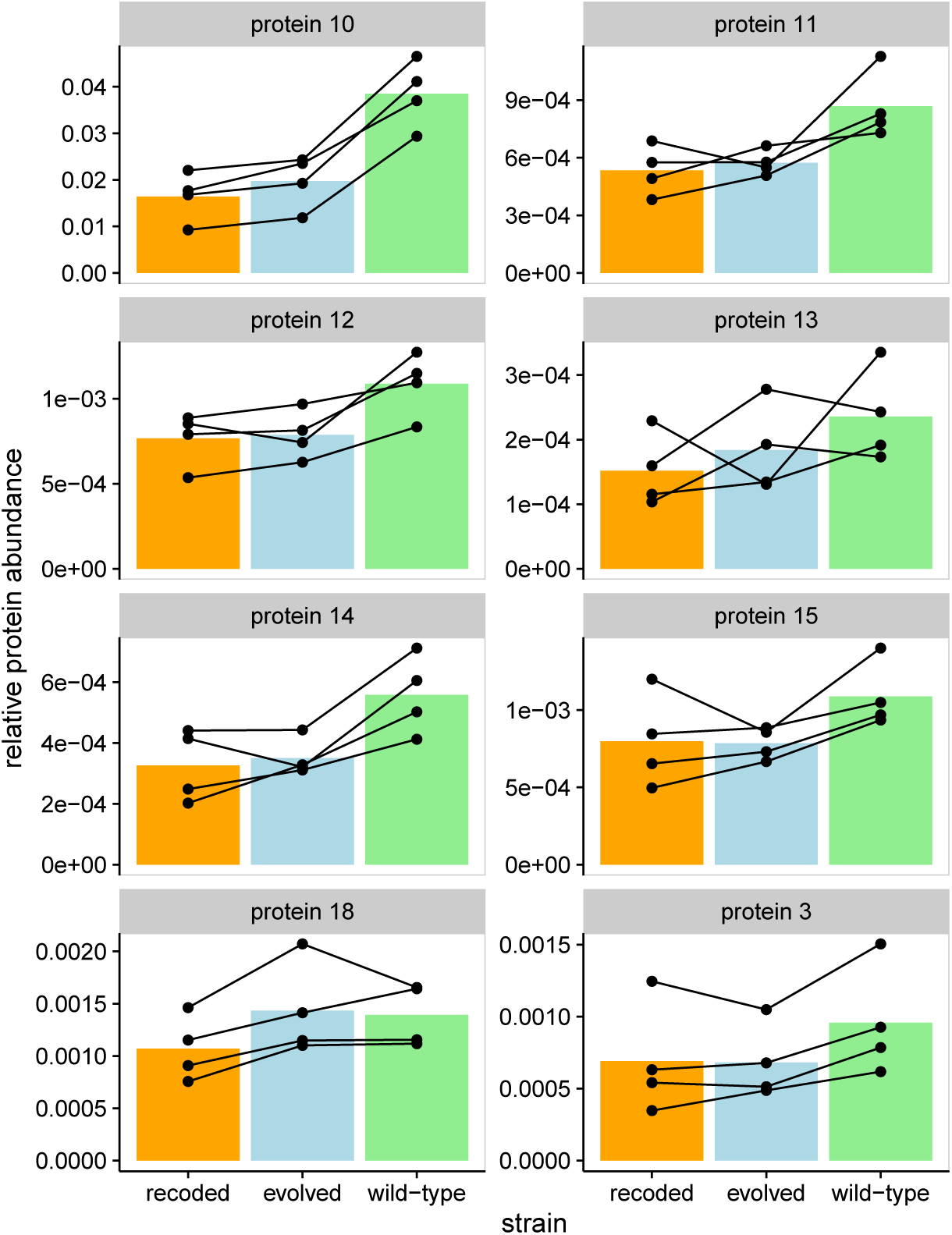
Recoding of gene *10A* reduces protein abundances of capsid protein, 5 other proteins encoded immediately downstream of *10A*, and the T7 endonuclease. The bacteriophage T7 genome contains many polycistronic transcripts. Genes *11* and *12* are only transcribed with gene *10*, following a read-through of the T*Ф* terminator. Genes *13*, *14*, *15* may also share the same transcript as gene 10, although these will be less common because of an RNAse cleavage site between genes *12* and *13*, and a promoter before gene *13*. We show the relative protein abundances corresponding to these genes, in addition to gene *3*, for all wild-type, recoded, and evolved strains at the 9 minute time point. All of these genes, with the exception of gene *3*, are class III genes, expressed late in the T7 life cycle. In addition to lower abundance of capsid protein (*10*), all protein products from the 5 genes immediately downstream of *10* are also suppressed (false discovery rate FDR < 0.1, FDR-corrected paired *t* test). These proteins are tail tubular proteins (*11* and *12*), probable virion-associated protein (*13*), and two internal virion proteins (*14* and *15*). Gene *3*, which codes for the T7 endonuclease, also has a reduced abundance in the recoded strain. Protein abundances for the evolved strain fall somewhere between that of the wild-type and the recoded strains. Each point represents a single measurement, and lines connect biological replicates.

Whereas proteins encoded downstream of the recoded gene *10A* showed decreased abundances, those encoded immediately upstream of gene *10A* were not obviously affected. Gene *9*, immediately upstream of *10*, encodes the highly expressed scaffold protein but this protein showed no difference in abundance between wild-type and recoded strains (Fig 4). Under the ribosome depletion hypothesis (model 2), we expected all genes expressed at the same time as gene *10* to be suppressed, thus including gene *9*. Indeed, because of the high levels of expression of *9* and the consequent ease of measuring it with our proteomics methods, we reject the ribosome depletion model as the basis of attenuation.

**Figure 4:**
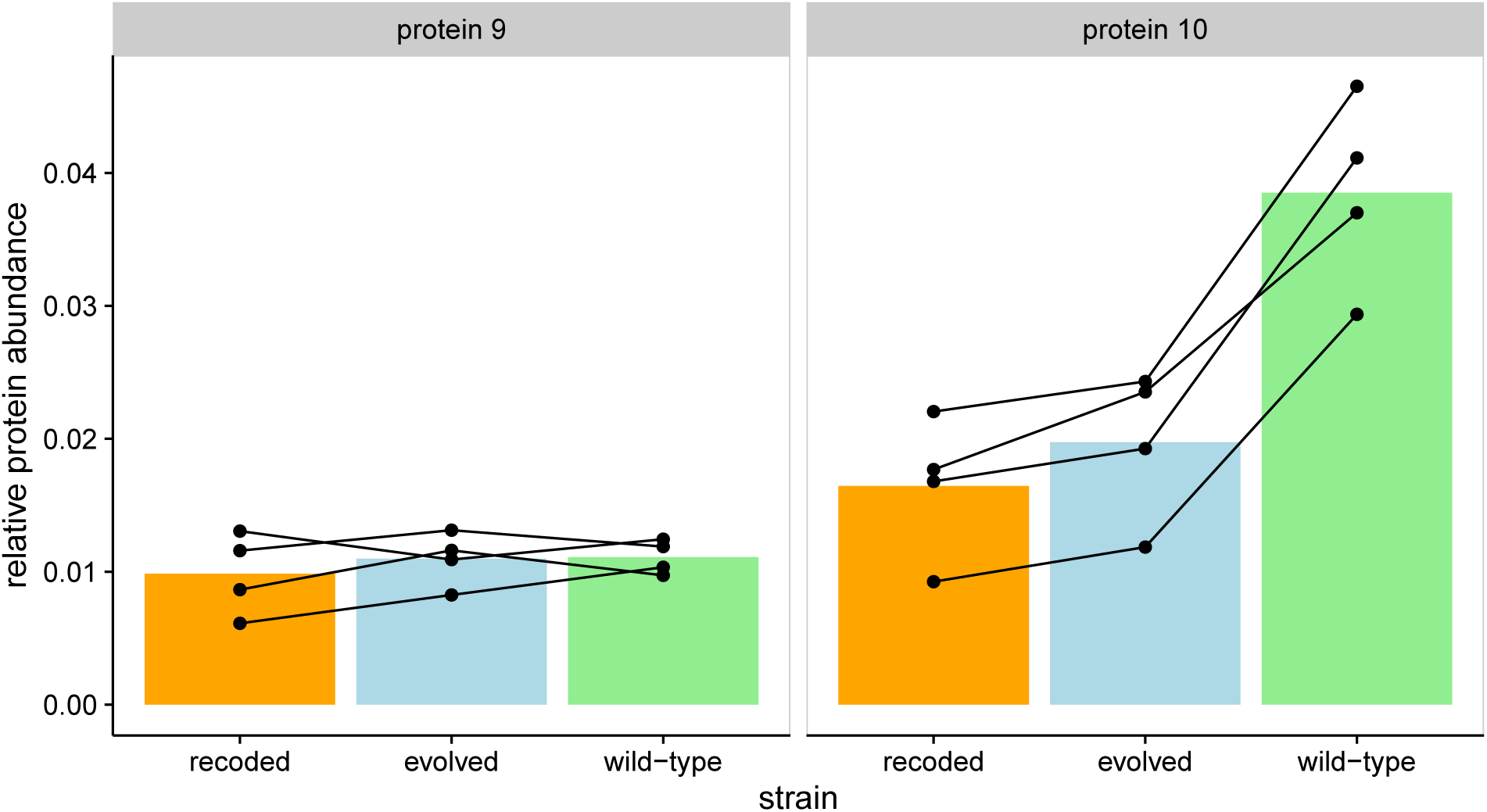
Gene *9*, encoding the highly-expressed scaffold protein, shows no detectable difference in abundance between wild-type and recoded strains. Genes *9* (left) and *10* (right) are both class III genes that are expressed at approximately the same time in the T7 life cycle. All abundances are from 9 minutes post-infection. Each point represents a single measurement, and lines connect biological replicates.

### E. coli mRNA transcripts declined after bacteriophage T7 infection

A reduction in capsid protein abundance in the attenuated virus could be explained by fewer transcripts from the gene. Of course, a reduction in transcription is not expected from a change in codon usage, and indeed, the design left the 5’-end of the gene unaltered specifically to keep transcription and translation initiation unchanged. Nonetheless, RNA-sequences were obtained from phage-infected *E. coli* at 1 minute, 5 minutes, and 9 minutes after infection to see if the decline in protein was accompanied by a decline in transcripts. Transcript abundances were normalized to the to total RNA abundance within a sample, excluding rRNA and tRNA. Both phage and *E. coli* transcripts were included.

Gene 10 transcript abundance increased over time, being highest at the final 9 minute time point. Yet no significant heterogeneity was observed among the wild-type, recoded, and evolved strain transcripts of gene 10 (Fig. 6A). This observation seems to rule out a transcriptional cause of capsid protein reduction.

As validation of the RNA-sequencing methods, transcription analyses were extended to other properties of the T7 and *E. coli* transcriptomes. Gene 1 (T7 RNA polymerase) transcript abundance decreased over time as a proportion of total transcripts (Fig 6B), consistent with previously observed class I gene expression timing (Molineux, 2005). Likewise, the relative abundance of T7 transcripts to *E. coli* increased sharply, consistent with established mechanisms by which T7 shuts off host transcription and degrades the host genome (Molineux, 2005). At 1 minute after infection, T7 transcripts comprised less than 1% of the transcript pool of infected *E. coli* (Fig 5). By 5 minutes after infection, T7 transcripts made up more than 75% of the transcript pool. By 9 minutes post-infection, this proportion reached approximately 95%. Although it is not possible to assess changes in absolute transcript abundances (see Materials and Methods), the data require some combination of host-cell transcripts being degraded rapidly, or T7 synthesizing transcripts so rapidly that they quickly dwarf the pool of *E. coli* transcripts. No strain-specific trends were detected.

**Figure 5:**
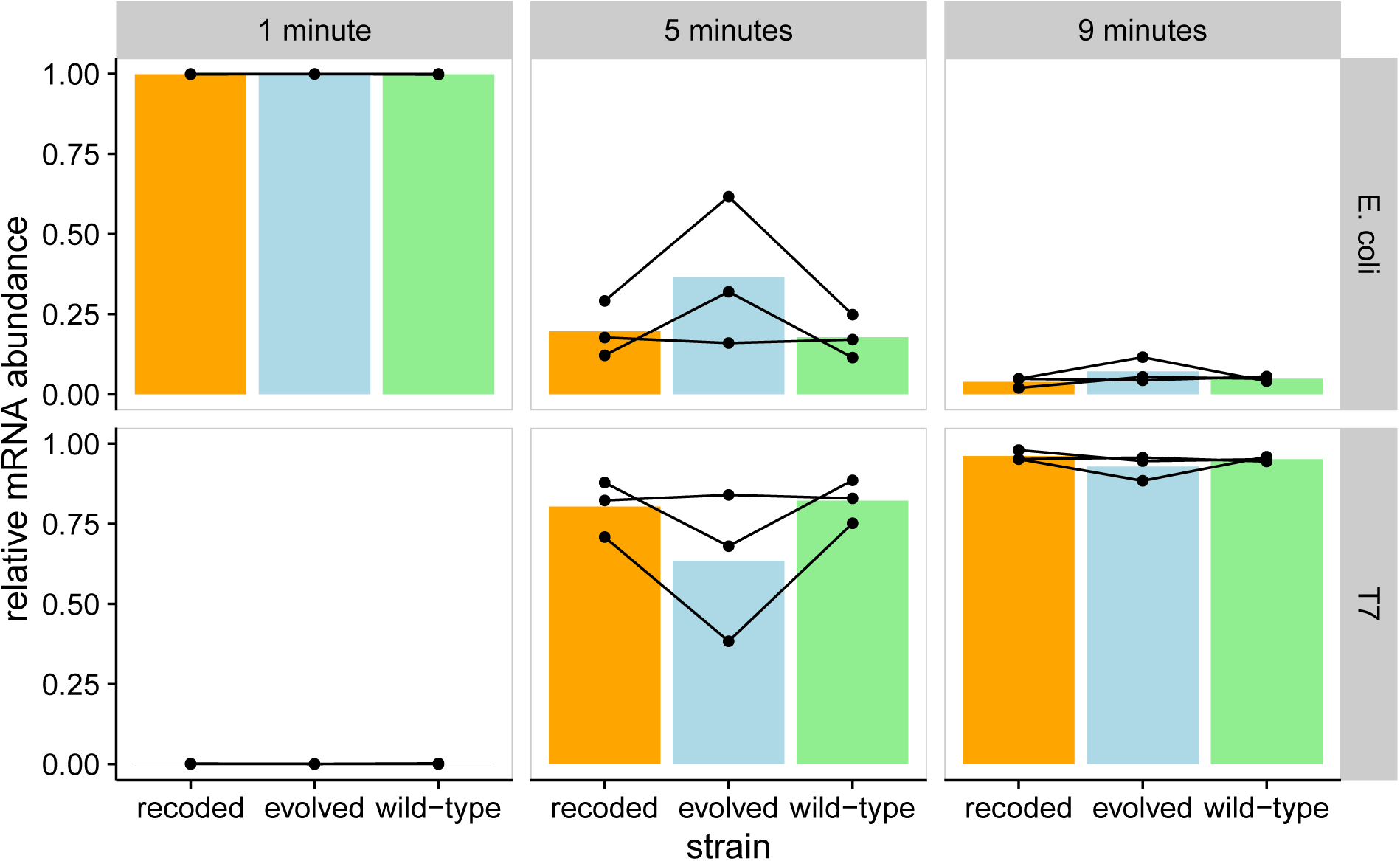
T7 transcripts make up most of the mRNA content of infected *E. coli*. The top row shows total *E. coli* transcript abundances, and the bottom row shows total T7 abundances. Points show individual experiments, while lines connect measurements from the same biological replicate. Each bar represents the mean relative mRNA abundance for a given strain and time point. Transcript abundances are shown relative to the total mRNA content of the sample. Wild-type, recoded, and evolved strains are shown, and there are no detectable differences between the three strains.

**Figure 6:**
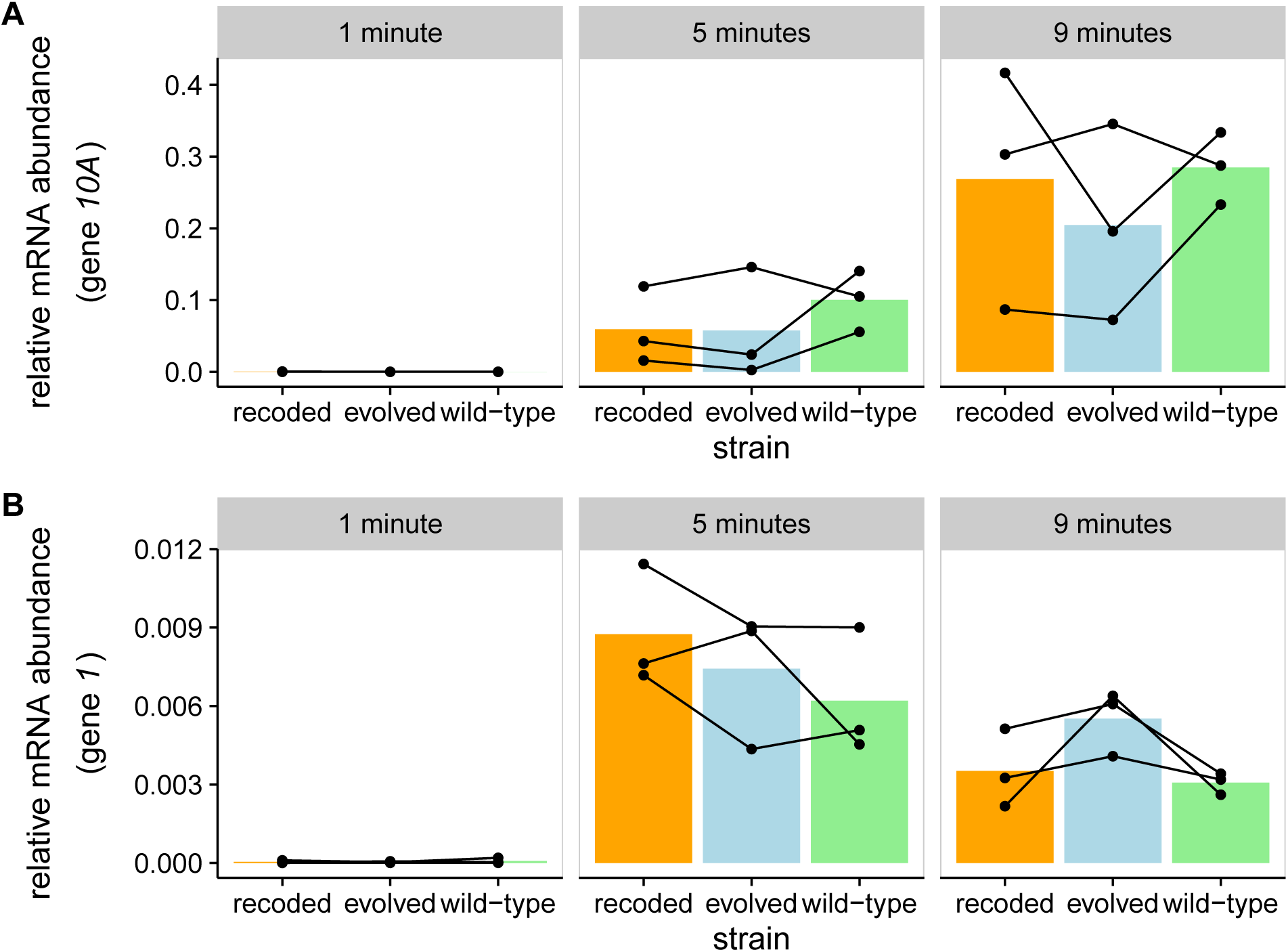
Transcript abundances of individual T7 genes change over time. Three biological replicates each are shown for wild-type, attenuated, and evolved strains. Transcript abundances are presented relative to the total mRNA content in the sample. Each column represents a different time point after infection. Both gene *10* and gene *1* follow the expected expression patterns of class I and class III genes, respectively. Each point represents a single measurement, and lines connect biological replicates. A Transcripts of gene *10* increase over time. There are no detectable differences in gene *10* mRNA abundances between the attenuated and wild-type T7 strains. B Transcripts of gene *1* increase from 1 to 5 minutes, then decrease from 5 to 9 minutes. Again, there are no detectable differences in gene *1* mRNA abundances between the attenuated and wild-type T7 strains.

### Downstream effects of recoding support a model of translational coupling

The proteomics suggest that translation of some downstream genes are specifically depressed by the recoding of *10*. This effect on downstream genes might involve translational coupling, in which a stalling of translation over *10* delays translation of genes further down on the same transcript. Translational coupling often occurs when multiple genes are encoded on a single transcript with little intergenic space, such as in bacterial operons and viruses (Lesage et al, 1992; Hellmuth et al, 1991; Schümperli et al, 1982; Oppenheim and Yanofsky, 1980; Aksoy et al, 1984; Torgov et al, 1998). Translational coupling is a plausible process for sets of T7 genes because most transcripts include multiple genes (Dunn and Studier, 1983). The T7 class III promoters precede genes 6.5 (the first class III gene), *9, 10, 13* and *17*; some transcripts with *9* and *10* will thus include earlier genes, but many will not (Fig 8). Although a phage-specific terminator between *10* and *11* aborts most (but clearly not all) *10* transcripts before *11*, all transcripts with *11* and *12* necessarily include *10*. Thus, translational coupling would operate for *11* and *12* if many of the ribosomes on those genes first translated *10*. Translational coupling beyond *12* is less plausible, however. The combination of an RNAse III site between *12* and *13* and a promoter before *13* will mean that many or most transcripts with *13* do not include *12*. So we expect substantially higher levels of translational coupling of *11* and *12* with *10*, but far less between *10* and *13*.

Polycistronic transcripts are necessary for translational coupling but not sufficient. It must also be the case that translation of downstream genes is usually from ribosomal re-initiation from the upstream genes, with few *de novo* ribosome initiations on the downstream genes. In turn, ribosome initiation can be impeded by secondary structure that sequesters the RBS (ribosome binding site) of the downstream genes (Tian and Salis, 2015; Rex et al, 1994; Qu et al, 2011). Ribosomes that reach the stop codon of one gene often expose the RBS of the next gene on the transcript and then reinitiate translation on that downstream gene (Spanjaard and van Duin, 1989). Support for the coupling model was evaluated from secondary structure predictions and in silico predictions of ribosome binding (Tian and Salis, 2015). Due to limitations in our software-based methods, we only tested the coupling of genes *10* and *11*. With translational coupling, translation initiation rates are predicted to be about 6 times greater for gene *11* than they would if gene *11* occurred on a single-gene transcript. This supports a model in which at least genes *10* and *11* are translationally coupled.

As the inference of translational coupling here is indirect and tentative, additional insight was sought from a mathematical model. The model assumed three genes on a single transcript in the order *a*, *b* and *c* (Fig 7A, eqs. 1–6 in Materials and Methods). Gene *b* is the one recoded to slow translation in some iterations of the model. Protein production rates (i.e. the rate at which ribosomes complete translation) depend on translation initiation and translation elongation rates. The model allowed varying degrees of coupling, with a coupling constant *q* assigned the values 0.2, 0.4, 0.6, and 0.8 (see Table 2 and Materials and Methods for all parameters). In a system with no coupling (*q* = 0), translation initiation of gene *b* does not depend on translation of gene *a*, and likewise between genes *b* and *c*. In a fully coupled system (*q* = 1.0), translation initiation of gene *b* depends entirely on the rate at which ribosomes complete translation of gene a. Translation of gene *c* similarly depends on gene b in a fully coupled system. We refer to this upstream-gene-dependent translation initiation rate as the effective initiation rate. Intermediate values of coupling are referred to as weakly coupled (*q* = 0.2), partially coupled (*q* = 0.4, 0.6), or strongly coupled (*q* = 0.8), and indicate that effective translation initiation rates depend on both elongation rates of upstream genes and rates of *de novo* initiation (indicated by the (1 − *q*) term in eqs. 2 and 3). To simulate codon deoptimization of one gene in the transcript, we varied the translation elongation rate of gene b only. All genes were assumed to be the same length. Thus, we explored how rates of protein production depend on both coupling between genes and elongation rates within genes.

**Table 2:**
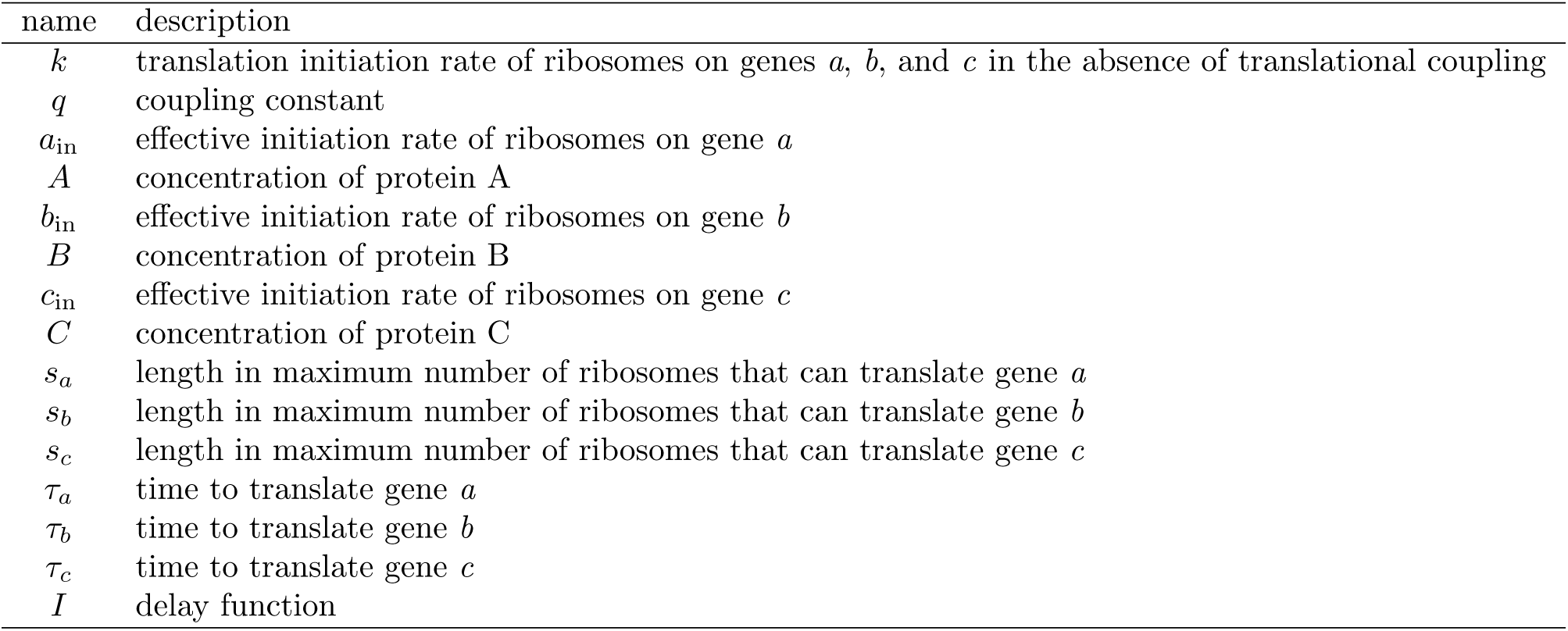
Parameters for 3-gene model of translational coupling.

| name | description |
| --- | --- |
| $k$ | translation initiation rate of ribosomes on genes $a$, $b$, and $c$ in the absence of translational coupling |
| $q$ | coupling constant |
| $a_{\text{in}}$ | effective initiation rate of ribosomes on gene $a$ |
| $A$ | concentration of protein A |
| $b_{\text{in}}$ | effective initiation rate of ribosomes on gene $b$ |
| $B$ | concentration of protein B |
| $c_{\text{in}}$ | effective initiation rate of ribosomes on gene $c$ |
| $C$ | concentration of protein C |
| $s_a$ | length in maximum number of ribosomes that can translate gene $a$ |
| $s_b$ | length in maximum number of ribosomes that can translate gene $b$ |
| $s_c$ | length in maximum number of ribosomes that can translate gene $c$ |
| $\tau_a$ | time to translate gene $a$ |
| $\tau_b$ | time to translate gene $b$ |
| $\tau_c$ | time to translate gene $c$ |
| $I$ | delay function |

This model demonstrated that recoding a gene in a translationally coupled set of genes affects the protein production rates of downstream, but not upstream, genes (Fig 7). Under strong translational coupling, the translation initiation rates of a gene depended entirely on the rate of ribosomes moving through the stop codon of an upstream gene. Thus the rate of protein A, B, and C production (assuming that elongation is slower than initiation in each gene) depended only on the translation initiation rate of gene a because *b* and *c* rely on re-initiation from *a*. In turn, when the translation time increased (i.e. elongation rate decreased) for gene *b*, translation elongation became rate-limiting and the production rates of proteins B and C decreased while A production rates remained unaffected (Fig 7B, right panel). Conversely, in a weakly coupled model, even if elongation rates decreased below initiation rate in gene *b*, production of protein C was only weakly affected by recoding of gene *b* (Fig 7B, left panel). In partially coupled models (Fig 7B, middle panels), the production rate of protein C also decreased as elongation rates decreased below initiation rates for gene *b*. However, this decrease in the rate of C protein production was smaller than the decrease observed on a strongly coupled model.

**Figure 7:**
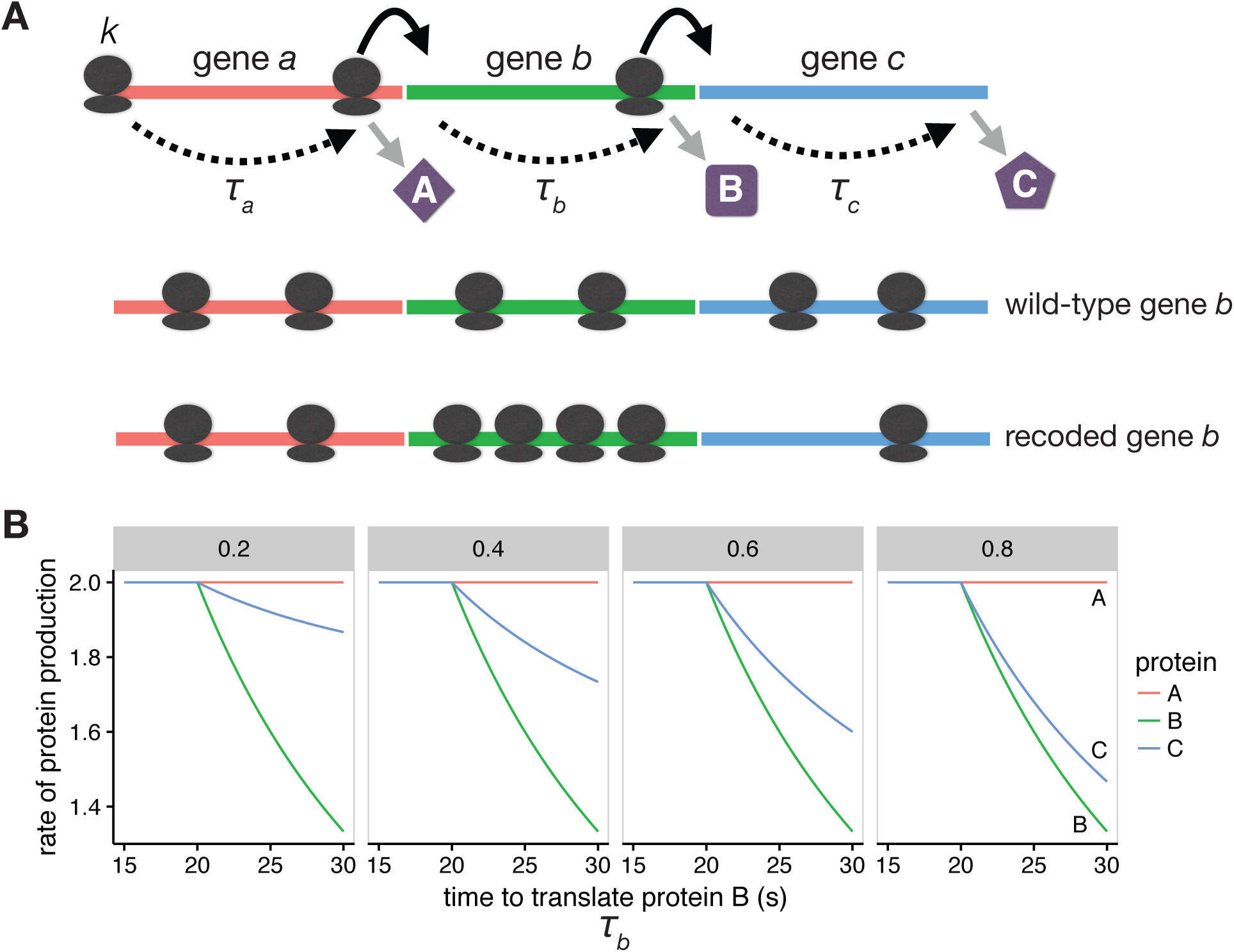
When genes are translationally coupled, the translation rate of upstream genes can affect translation rates of downstream genes. A Here we show three genes (*a*, *b*, and *c*) in a polycistronic transcript that are translationally coupled. That is, we assume that the translation initiation rates of one gene depend partially on the rate of ribosomes reaching the stop codon of the previous gene. Typically, translation initiation is the rate-limiting step in translation. All genes are assumed to be the same length. Translation initiation of gene a is given as *k*, and the translation time (after initiation) is given as *τ*. Upon recoding gene *b*, we hypothesize that the ribosomal density on *b* increases (bottom row). B As translation elongation rates decrease below initiation rates, the rate of translation of downstream genes also decreases. The translation rate of *b* and *c* decline sharply as the translation elongation rate of gene *b* decreases below the initiation rates of *b* and *c*. The left, middle, and right panels show models with coupling constants (*q*) of 0.2, 0.4, 0.6, and 0.8 respectively. A larger coupling constant indicates that downstream genes are less likely initiate translation *de novo* and instead rely on ribosomes from upstream genes. As coupling increases, production of protein C becomes more sensitive to translation elongation rates of gene *b*.

### Connecting proteomics to fitness

In previous work, the T7 with recoded capsid gene had been found to have a fitness of 35.7 doublings/hr, compared to a value of 43.2 for the wild type. Here we consider whether and how the altered proteomics might lead to this fitness reduction. The connection from proteomics to fitness spans two steps: (i) identify the phage life-history components affected by the recoding and evaluate whether that change is compatible with the proteomics, then (ii) assess whether the magnitude of altered fitness components is compatible with overall fitness.

In the growth conditions used for our assays, fitness is determined by cell density and three phage properties: burst size, lysis time, and adsorption rate (Wang et al, 1996; Shao and Wang, 2008; Guyader and Burch, 2008; Bull, 2006; Patwa and Wahl, 2009; Bull et al, 2011). Burst sizes and lysis times were estimated here for the wild-type and recoded phages (there was no expectation that adsorption rate would be affected, which depends on the presence of tail fibers, the product of gene *17*). Although the same cell line was used here as in previous studies (IJ1133), new cell preparations were used, so quantitative agreement with past estimates of burst size and lysis time is not expected, but proportional differences should scale across different cell preparations. No difference in lysis time was observed between strains, but burst size was reduced almost 50% with the recoding (Fig 9).

**Figure 8:**
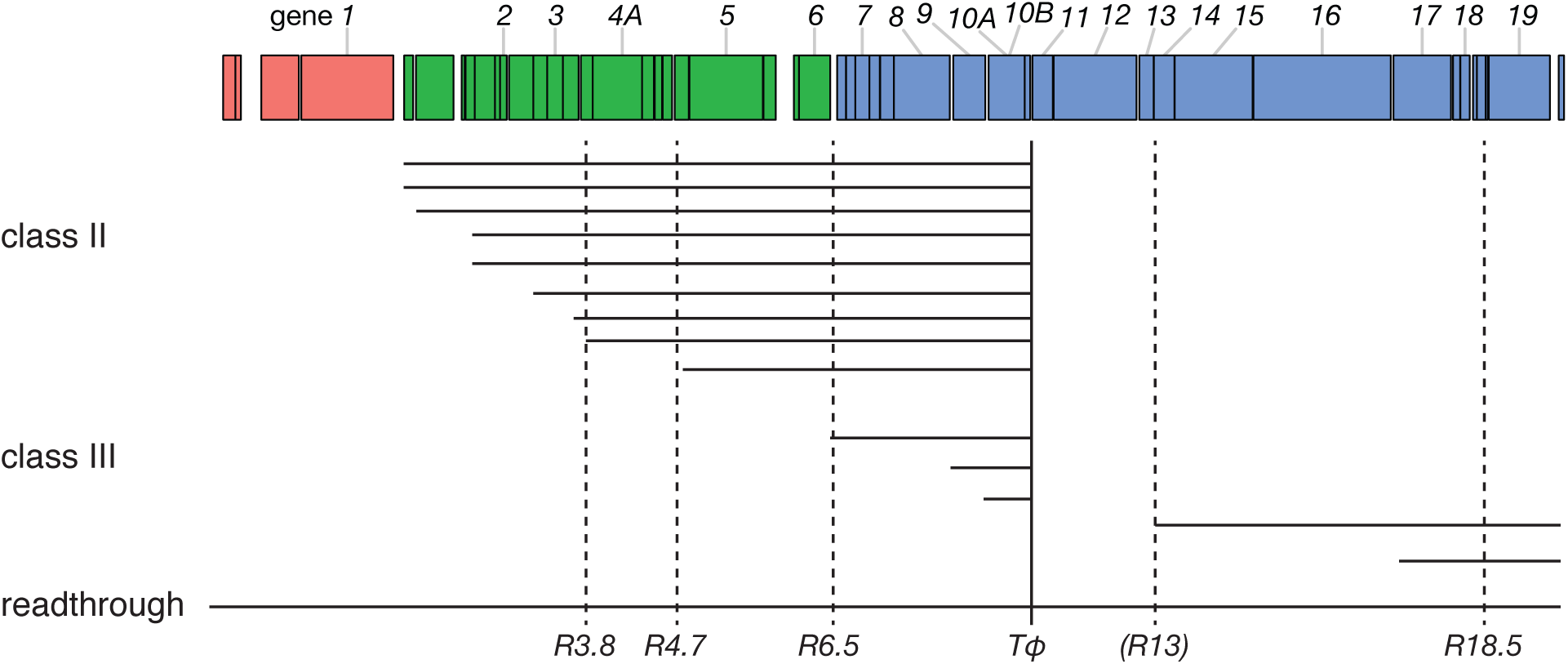
T7 produces many polycistronic transcripts. The bar across the top shows the T7 genome, and each class is shown in a different color. Horizontal lines represent transcripts. Dashed vertical lines represent RNAse cleavage sites, where R*3.8*, R*4.7*, R*6.5*, and R*18.5* are strong cleavage sites. R*13* is a weak RNAse cleavage site. The solid vertical line represents the terminator T*ø*. Genes *11* and *12* are only ever expressed as a product of read-through of T*ø*, indicated by the read-through transcript. Only transcripts containing class III genes are shown. Not all read-through products are shown. Data are from Dunn and Studier (1983).

**Figure 9:**
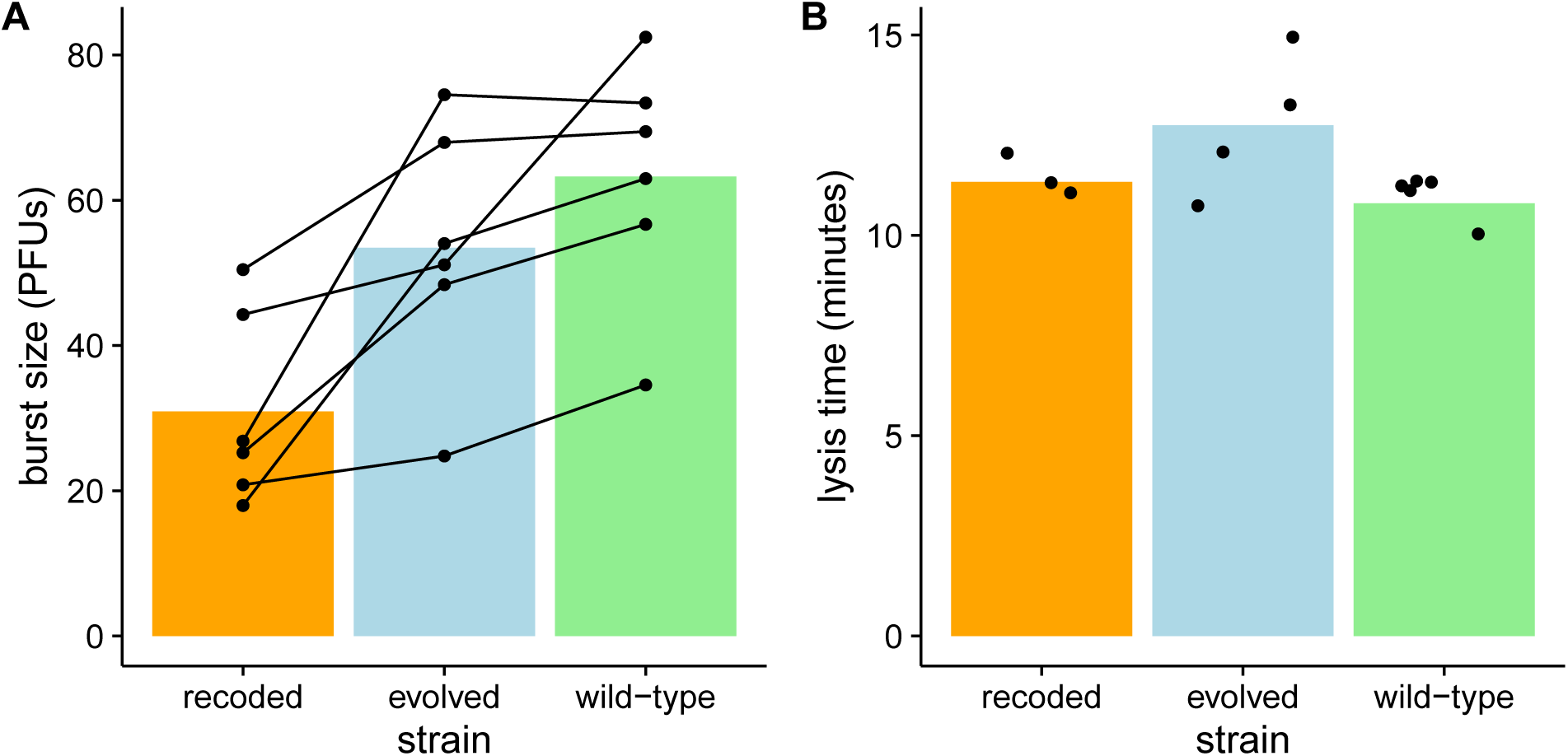
Burst size in recoded T7 strain is lower than that of the wild-type and evolved strains, while lysis time is indistinguishable for all three strains. A Five replicates each of burst size measurements are shown for the wild-type, recoded, and evolved T7 strains. Points represent individual measurements in plaque forming units (PFUs), and each set of measurements in a replicate are connected by a line. Bars show the mean burst size for each strain. Burst size of the recoded strain is smaller than that of the evolved (*p* = 0.02, paired *t* test) and wildtype (*p* = 0.002, paired *t* test) strains. B Lysis time for the three different strains is shown. Points represent individual measurements, and bars represent the mean lysis time for each strain. There were no significant differences between the lysis times of each strain (*p* = 0.4, ANOVA).

The proportional reduction in burst size is nearly the same as that for the reduction in capsid protein abundance at 9 minutes. This reduction in burst size is no doubt caused by the reduction in capsid protein. There is perhaps little basis for arguing that the reductions should match quantitatively, but the agreement between the two numbers poses no dilemma.

The second step in connecting proteomics with fitness is to consider whether a 50% reduction in burst size (with no change in lysis time) is compatible with the observed fitness reduction of 7.5 doublings/hr. The mapping of phage fitness components onto total fitness has been addressed in detail (Bull et al, 2011). From numerical trials in that study, a 50% reduction in burst size is compatible with a fitness reduction of the magnitude observed here (their Table 1, lines L1 and L5).

## Discussion

In the decade since the first proposals to attenuate viruses by synonymous codon substitutions, it has been established that the method works in many viruses and offers many advantages over earlier methods of attenuation. Yet the mechanism by which silent codon changes attenuate not only remains elusive but seems less clear now than it did at the start. Nor is it clear that a single mechanism underlies the attenuation in different systems. Here, we extended previous work on a bacteriophage system in which the encoding of rare codons in the major capsid gene reduced fitness. Our goal was to refine an understanding of the molecular basis of the attenuation.

The capsid protein (encoded by gene *10*) is the most abundant protein produced during the infection cycle of bacteriophage T7. Deoptimizing 50% of gene 10 codons reduced fitness (Bull et al, 2012). In exploring the underlying molecular mechanism by which the recoding has this effect, our primary result is that the protein product of gene 10 is reduced almost 50% by the end of the infection cycle, but protein abundance of genes immediately downstream of gene 10 are also depressed. The differences in protein abundance are not reflected in transcript levels, so it appears that the suppression of protein levels lies in translation. The evidence thus supports a simple interpretation of the fitness impact of recoding the major capsid gene:

1. Capsid protein is expressed at a reduced level, as are a few downstream genes.
2. Burst size is correspondingly reduced approximately 50% with no change in lysis time, compatible with the observed reduction in total fitness.

One mechanism we entertained to explain the altered proteomics of the recoded phage is saturation of the ribosomes with gene *10* transcripts. Such a model requires that the production of all T7 proteins would decline late in the infection cycle for the recoded phage. Within the limits of resolution, the proteomics rule out an overall reduction in T7 protein production, indicating that reductions are limited to the recoded *10* and a few downstream genes. This study may provide the first indication that translational effects of the recoding extend beyond the recoded genes. There was also no evidence for aborted gene *10* polypeptides in the recoded strains, as might occur from ribosomes stalled on gene *10* transcripts. An obvious next step is to extend these analyses to ribosome profiling, which would directly indicate whether the recoding does tie up ribosomes on gene *10* (Li et al, 2014).

The means by which synonymous codon replacement attenuates may be more straightforward for phage T7 than for eukaryotic viruses. Several lines of evidence suggest that the eukaryotic virus attenuation by synonymous codon changes is from the creation of CpG dinucleotides (Burns et al, 2009; Atkinson et al, 2014; Tulloch et al, 2014); indeed, especially powerful evidence to support this interpretation is that evolutionary reversions of attenuated viruses disproportionately reverse CpGs (Burns et al, 2006). In contrast, T7 evolutionary reversions did not exhibit any signature suggestive of a dinucleotide basis for attenuation (Bull et al, 2012). Nonetheless, we expect that the mechanism of attenuation in T7 will apply across other viruses, even if it is not the only mechanism operating in those viruses.

If our interpretation is correct for the mechanism underlying the effect of the recoding, an evolutionary response to overcome the effect might be duplication of promoter immediately upstream of gene *10*. Such a duplication would increase the number of *10* transcripts and, in the absence of ribosome saturation, would increase the amount of capsid protein. A duplication of a class II promoter was observed during adaptation of a different phage (Springman et al, 2005), adding credence to the possibility of such an outcome. Yet the mutational origin of a promoter duplication may be highly dependent on surrounding sequences so perhaps not feasible for all promoters.

We propose that translational coupling may explain why expression of genes downstream of *10* is suppressed by the recoding. Part of that inference is based on a mathematical model of translation. That model is necessarily simplifed, however, and there are some obvious improvements needed to increase its realism. First, it assumes a fixed quantity of transcripts, when we know from our RNA-sequencing results that T7 transcripts increase rapidly during infection. Second, the model assumes a per-gene translation elongation rate, but does not model individual codons. A more sophisticated model that includes codon-level detail and the T7 life cycle would be needed to predict the fitness effects of codon deoptimization. Several life-cycle and molecular models of T7 have achieved limited success in predicting the phenotypic effects of genome manipulations (Endy et al, 1997; You et al, 2002; Kosuri et al, 2007; Birch et al, 2012), but none enable codon modifications or complex translation mechanisms such as coupling. The proteomics and RNA-sequencing data generated in this study should be useful in future high-resolution modeling studies that scale from the molecular level to that of viral fitness.

Although it is tempting to interpret the slowed translation as a consequence of using rare codons, which then use rare tRNAs, some recoding strategies used in other genomes suggest alternative possibilities. Codon deoptimization of GFP in *E. coli* initially yielded a range of protein expression effects, but these effects were eventually attributed to changes in mRNA secondary structure in the first 28 codons of the GFP sequence (Kudla et al, 2009). Codon changes beyond these first 28 codons had a weak effect on protein expression (Kudla et al, 2009). In our designs, we explicitly excluded these 5’-end codons from modification. Moreover, some attenuation designs with influenza virus and poliovirus have achieved attenuation by merely shuffling codons within a gene or genome to create rare codon pairs. Since the numbers of each codon are not changed in those designs, the mechanism cannot be one of simple tRNA abundance. One possibility is suggested by recent work in yeast, whereby the degree of codon clustering is important to rapid translation (Cannarozzi et al, 2010; Gamble et al, 2016). We did not attempt to control for codon-pair bias in our recoded T7 constructs. Further experiments will be needed to determine if prokaryotic translation machinery, like that of the eukaryotes, is sensitive to changes in codon-pair bias.

The approach developed here should help elucidate other mechanisms of viral attenuation. For example, the timing of gene expression appears important to fitness: a reciprocal exchange of some middle and late genes had some major fitness effects, and those effects were not recovered on long term adaptation (Cecchini et al, 2013). Ultimately, we envision a future in which an understanding of viral life history at the molecular level enables facile engineering of arbitrary fitness and alternative vaccine designs.

## Materials and Methods

### Gene *10* nomenclature

Gene *10* is translated in two forms, A and B. Form A is 344 amino acids and is formally denoted the major capsid protein. Form B (the minor capsid protein) is not essential and results from a ribosomal frameshift at the end of A and is 397 amino acids. In the engineering, all codon changes were within *10A* and thus also within *10B*. Moreover, since most peptide fragments coming from the minor and major capsid proteins ambiguously mapped to both proteins, abundances of 10A and 10B were not differentiated using our proteomic methods and were combined. We followed a similar procedure for our RNA-sequencing analyses. To simplify notation, we merely refer to the recoded gene as *10* and the affected A and B proteins as capsid protein.

### Bacteriophage T7 strains and *E. coli* hosts

The host for all experiments was IJ1133 [*Escherichia coli* K-12, F-ΔlacX74 thiΔ (mcrC-mrr)102::Tn10]. T7 strains used in this study come from (Bull et al, 2012). An isolate of T7_61_ (a population adapted to grow optimally on IJ1133 specifically through serial passage) was first deleted of its gene *10A*, then recombined over a plasmid carrying a different gene *10A* engineered to contain a low fraction (0.1) of preferred codons. The recombinant, denoted here as the recoded strain, could be identified by its ability to grow without complementation. The evolved strain was initiated from the recoded phage and adapted over 800–1000 generations [strain L1 from (Bull et al, 2012)]. The wild-type strain in this study was derived from the recoded strain after recombination over a plasmid containing wild-type gene 10, then grown out for 6 hours of serial transfer on IJ1133. Fitness of this ‘wild-type’ strain was approximately the same as that of the ancestral population (T7_61_).

### Burst size and lysis time

Lysis time and burst size assays were performed as previously described (Heineman and Bull, 2007; Bull et al, 2011). The initial infection steps were identical for both assays. Briefly, 10^8^ phage (MOI = 0.1) were added to a 10 mL culture of exponentially growing cells (37°C with agitation), incubated for 3 minutes and subsequently diluted 10^4^-fold to prevent further adsorption. For lysis times, phage were plated at various time points between 4 and 18 minutes (after initial infection) to monitor changes in titer; lysis time was taken as the time of the first significant increase in titer.

To determine burst size, initial density of phage-infected cells was determined by plating phage before and after treatment with chloroform 5 and 6 minutes after initial infection. Cells infected with phage at the time of chloroform treatment do not produce viable phage, so only free phage will form plaques, allowing for the determination of phage-infected cells at these times. Final phage titers were obtained at 15, 16, and 17 minutes by plating chloroform-treated samples. Burst size was then calculated as the phage titer at the end time points divided by the number of initial phage-infected cells.

### RNA sequencing

*E. coli* was grown in LB broth to a concentration of 10^8^ cells/mL at 37°C with agitation, then infected with phage at an MOI of 2.5. At 1, 5, and 9 minutes post-infection, 2 mL of bacterial suspension were removed from the phage-infected cultures and pelleted in a microcentrifuge. Pellets were either flash frozen in liquid nitrogen or immediately used for downstream processes (RNA extraction or protein preparation for proteomics). RNA was isolated using Trizol reagent, following the manufacturer’s protocol. Library preparation and sequencing was performed by the University of Texas Genome Sequencing and Analysis Facility (GSAF) using Illumina NextSeq 500 (SR75).

Since gene *10B* is a readthrough product of gene *10A*, we excluded the gene *10B* transcript from the reference transcriptome. RNA-sequencing reads were quantified using Kallisto (Bray et al, 2016) and *E. coli* K-12 (NCBI: U00096.3) and T7 (NCBI: NC_001604.1) reference genomes. For analyses of the recoded and evolved T7 strains, the gene *10A* sequence was replaced with the recoded sequence (Bull et al, 2012, Supplementary file S1) in the reference genome. A population of T7-infected *E. coli* has no core set of stably-expressed genes with which to normalize during differential expression analysis. Therefore, all transcript abundance estimates (transcripts per million, TPM) were normalized to the total cellular transcript abundance (including both T7 and *E. coli* transcripts). Differential expression analysis was only possible within genes, but not between them.

### Proteomics

Proteomics was performed as previously described in Houser et al (2015). In brief, T7-infected *E. coli* cell pellets (prepared as described in the RNA sequencing section above) were resuspended in 50 mM Tris-HCl pH 8.0, 10 mM DTT. 2,2,2-trifluoroethanol (Sigma) was added to 50% (v/v) final concentration and samples were incubated at 56°C for 45 minutes. Following incubation, iodoacetamide was added to a concentration of 25 mM and samples were incubated at room temperature in the dark for 30 minutes. Samples were diluted 10-fold with 2 mM CaCl_2_, 50 mM Tris-HCl, pH 8.0. Samples were digested with trypsin (Pierce) at 37°C for 5 hours. Digestion was quenched by adding formic acid to 1% (v/v). Tryptic peptides were bound, washed, and eluted from HyperSep C18 SpinTips (Thermo Scientific). Eluted peptides were dried by speed-vac and resuspended in Buffer C (5% acetonitrile, 0.1% formic acid) for analysis by LC-MS/MS.

For LC-MS/MS analysis, peptides were subjected to separation by C18 reverse phase chromatography on a Dionex Ultimate 3000 RSLCnano UHPLC system (Thermo Scientific). Peptides were loaded onto an Acclaim C18 PepMap RSLC column (Dionex; Thermo Scientific) and eluted using a 5–40% acetonitrile gradient over 250 minutes at 300 nL/min flow rate. Eluted peptides were directly injected into an Orbitrap Elite mass spectrometer (Thermo Scientific) by nano-electrospray and subject to data-dependent tandem mass spectrometry, with full precursor ion scans (MS1) collected at 60,0000 resolution. Monoisotopic precursor selection and charge-state screening were enabled, with ions of charge > +1 selected for collision-induced dissociation (CID). Up to 20 fragmentation scans (MS2) were collected per MS1. Dynamic exclusion was active with 45 s exclusion for ions selected twice within a 30 s window.

We assigned each peptide to a protein or protein group (in the case of ambiguous peptides which map to multiple proteins) using Proteome Discoverer (Thermo Scientific) and REL606 and T7 reference proteomes (NCBI: NC_012967, NC_001604.1) concatenated with a database of contaminant proteins (http://www.biochem.mpg.de/5111795/maxquant). We selected the top three most abundant peptides by peak area for each protein. We averaged the these peptide peak areas across technical replicates to obtain a protein abundance estimate (Silva et al, 2006). All protein abundance estimates were normalized to the total *E. coli* protein content of the sample.

To determine if gene *10* C-terminal peptides were more common than N-terminal peptides in the recoded T7 strain compared to the wild-type strain, we compared the fit of the following two models [given here in notation from the lme4 (Bates et al, 2015) R package]: 

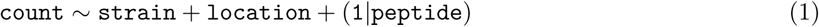

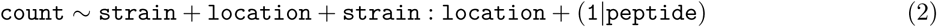

where `count` is the number of peptides of type `peptide, strain` is the strain of T7, and `location` is the location of `peptide` within gene *10*. We assume `peptide` to be a random effect, given by the (1 | peptide) term. The term `strain:location` indicates interaction between `strain` and `location`. Thus, we compare a model in which the location of a peptide and the strain interact, and a model in which there is no such interaction. If N-terminal peptides were less prevalent than C-terminal peptides in the recoded T7 strain, we would expect equation 2 to provide a better fit than equation 1.

## Models of translational coupling

### Biophysical model

Secondary structure near the ribosome binding site can inhibit translation initiation. On polycistronic transcripts, this secondary structure can be disrupted by ribosomes completing translation of an upstream gene, thus increasing translation initiation rates. Estimates of this relative increase in translation initiation of gene *11* due to translational of gene *10* and were predicted using the Operon Calculator (Tian and Salis, 2015).

### Mathematical model

To model the effects of translational coupling on rates of protein production, we first assume that we have a polycistronic transcript with genes *a*, *b*, and *c*. For the effective translation initiation rate *a*_in_ of gene *a*, we define the following relationship 

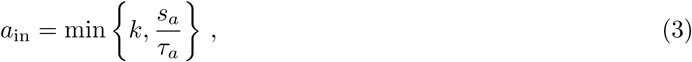

where *k* is some arbitrary initiation rate, *s_a_* is the length of gene *a* and *τ_a_* is the time it takes to translate gene *a*. As the time to translate gene *a* increases, the elongation rate will eventually be less than *k* and become the rate-limiting step as the transcripts back up with ribosomes. For the effective initiation rate *b*_in_ of gene *b* we define the following relationship 

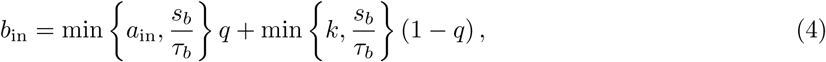

where *s_b_* and *τ_b_* are the length of gene *b* and the time to translate gene *b*, respectively. The constant *q* indicates the amount of translational coupling (the proportion of ribosomes terminating translation on gene *a* and reinitiating translation on gene *b*), where 0 is no coupling and 1 is fully-coupled. The term (1 − *q*) indicates the proportion of translation initiation events occurring *de novo*. Thus, the effective initiation rate is the sum of both the reinitiation rate from gene *a* to gene *b*, and *de novo* initiation rate of gene *b*. Similarly, the effective initiation rate *c*_in_ of gene *c* is 

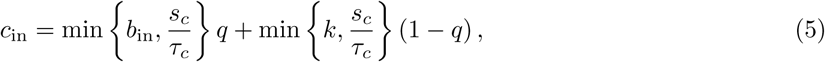

where *s_b_* and *τ_b_* are the length of gene *c* and the time to translate gene *c*, respectively. The coupling constant *q* and initiation rate *k* are the same as defined previously.

Now that we have defined effective initiation rates for genes *a*, *b*, and *c*, then the protein production rates 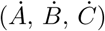 are defined as 

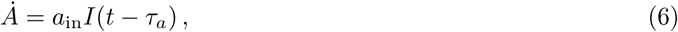

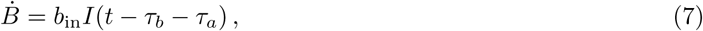

and 

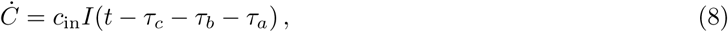

where each rate depends on the effective translation initiation rate and a delay function *I*(*t*). Lastly, we define the delay function as 

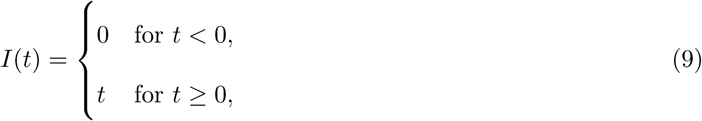

where *t* is time.

#### Statistical software and plots

All statistical tests were conducted using the R language (R Core Team, 2014). All plots were generated using the ggplot2 package (Wickham, 2009).

#### Data availability

We are in the process of submitting raw RNA reads to NCBI GEO (Barrett et al, 2013) and raw MS proteomics data to ProteomeXchange (Vizcaíno et al, 2014). All processed data and scripts are available at: http://github.com/benjaminjack/t7_attenuation.

## Acknowledgements

We thank I. J. Molineux for advice on and insight into T7. This work was supported in part by National Institutes of Health Grants R01 GM088344, National Science Foundation Cooperative agreement no. DBI-0939454 (BEACON Center), and Army Research Office Grant W911NF-12-1-0390. The Texas Advanced Computing Center provided high-performance computing resources. JBB is supported as the University of Texas Miescher Regents Professor.

### Author contributions

Conceived and designed the experiments: JJB COW DRB. Performed the experiments: DRB MLP BLS. Analyzed the data: BRJ DRB MLP BLS. Developed model: BRJ COW JJB. Provided reagents and materials: JJB COW DRB. Wrote the paper: JJB COW DRB MLP BLS BRJ.

### Conflicts of interest

The authors report no conflicts of interest.

